# Expression of fatty acid transport protein in retinal pigment cells promotes lipid droplet expansion and photoreceptor homeostasis

**DOI:** 10.1101/287672

**Authors:** Daan M Van Den Brink, Aurélie Cubizolle, Gilles Chatelain, Francesco Napoletano, Pierre Dourlen, Nathalie Davoust, Laurent Guillou, Claire Angebault-Prouteau, Nathalie Bernoud-Hubac, Michel Guichardant, Philippe Brabet, Bertrand Mollereau

## Abstract

Increasing evidence suggests that dysregulation of lipid metabolism is associated with neurodegeneration in retinal diseases such as age-related macular degeneration and in brain disorders such as Alzheimer’s and Parkinson’s diseases. Lipid storage organelles (lipid droplets, LDs), accumulate in many cell types in response to stress, and it is now clear that LDs function not only as lipid stores but also as dynamic regulators of the stress response. However, whether these LD functions are always protective or can also be deleterious to the cell is unknown. Here, we investigated the consequences of LD accumulation on retinal cell homeostasis in transgenic flies and mice overexpressing fatty acid transport protein (FATP) in retinal pigment cells (RPCs). In wild-type *Drosophila*, overexpression of *dFatp* specifically in RPCs resulted in an expansion of LD size in both RPCs and in neighboring photoreceptors but was non-toxic. Similarly, in mice, LD accumulation induced by RPC-specific expression of human *FATP1* was non-toxic and promoted mitochondrial energy metabolism in both RPCs and photoreceptor cells. In contrast, RPC-specific *dFatp* knockdown reduced neurodegeneration in *Aats-met^FB^ Drosophila* mutants, which carry a defective respiratory chain, indicating that abnormal LD accumulation can be toxic under pathological conditions. Collectively, these findings indicate that FATP-mediated LD formation in RPCs induces a non-autonomous increase of LDs in photoreceptors that promotes homeostasis under physiological conditions but can be deleterious under pathological conditions.

**Author Summary:** Lipids are major cell constituents and are present in the membranes, as free lipids in the cytoplasm, or stored in vesicles called lipid droplets (LDs). Under conditions of stress, lipids stored in LDs can be released to serve as substrates for energy metabolism by mitochondria. However, lipid storage is deregulated in many degenerative disorders such as age-related macular degeneration and Alzheimer’s disease. Thus, it is unclear whether accumulation of LDs is protective or can also be toxic. To address this question, we examined the consequences of enforced LD accumulation on the health of retinal cells in flies and mice. Like humans, fly and mouse retinas contain retinal pigment cells (RPC) that support the functions of neighboring photoreceptor cells. We found that overexpression of the fatty acid transport protein (FATP) in RPCs induced accumulation of LDs in both transgenic flies and mice. Moreover, LD accumulation in RPCs had a beneficial effect on juxtaposed photoreceptors under normal physiological conditions, but was toxic under pathological stress conditions. We propose that lipid storage is a mechanism of cellular communication that is essential to maintain photoreceptor health.

## Introduction

Photoreceptor neurons are among the highest energy consumers in the body. They are sustained by a layer of retinal pigment epithelial cells (here termed retinal pigment cells [RPC]) that provide photoreceptors with a constant supply of substrates for energy production by mitochondrial oxidation via the tricarboxylic acid (TCA) cycle (S1 Fig.). An inability of RPCs to perform this role leads to photoreceptor death and is associated with diseases such as age-related macular degeneration (AMD), the most prevalent cause of blindness in developed countries [1].

Injury to RPCs, whether through disease or as part of the normal aging process, is accompanied by an accumulation of lipid deposits, named drusen, in the RPC itself or in the adjacent Bruch’s membrane [2–4]. A common cause of injury is reduced blood flow to the retina, which causes hypoxia and a subsequent metabolic shift that results in accumulation in RPCs of intracellular lipid droplets (LDs) composed of neutral lipids, such as triacylglycerides (TAGs) and sterol esters. In AMD, lipids are also deposited extracellularly, further compromising the functions of RPCs and photoreceptors [5]. However, it is not clear whether lipid accumulation is a cause or a consequence of AMD pathology.

Work in *Drosophila* models has contributed greatly to our understanding of the mechanisms of LD biosynthesis and physiological functions [6, 7]. For example, LDs have been shown to play an antioxidant role in the developing *Drosophila* nervous system by protecting polyunsaturated fatty acids from peroxidation under conditions of hypoxia [8]. In contrast, in *Drosophila* carrying mutations affecting the mitochondrial respiratory chain, oxidative stress induces glial cells to accumulate LDs that are toxic to the adjacent photoreceptor cells [9]. Thus, LDs are not simply cytoplasmic lipid storage organelles with critical roles in energy metabolism; they also have dynamic functions in regulating the response to stress of many cell types, including those in the nervous system [10–12].

LDs are synthesized on the endoplasmic reticulum membrane by a protein complex composed of diacylglycerol acyltransferase (DGAT-2) and fatty acid transport protein (FATP) [12, 13]. The FATP family (also known as solute carrier family 27 [SLC27]) [14, 15] are acyl-CoA synthetases involved in the cellular import of Fatty Acids (FAs) [16] as well as other processes. FATPs transfer acyl-CoA to DGAT-2 for the synthesis of TAGs, which are then incorporated into expanding LDs. An important role for FATP in TAG storage has also been demonstrated in mammals. In *Fatp1* knockout mice, the uptake of FA and the size of LDs in the brown adipose tissue are reduced compared with wild-type (WT) mice, resulting in defects in nonshivering thermogenesis [17]. The FATP1 gene is conserved from yeast to humans, where there are six proteins with different tissue distributions and substrate preferences. *Drosophila Fatp* (*dFatp*) shows a high degree of sequence similarity to human (*h*)*FATP1* and *hFATP4*, which have broad expression profiles that include the brain and retina [16, 18, 19].

The *Drosophila* retina is composed of approximately 800 ommatidia, each of which comprises eight photoreceptors surrounded by a hexagonal lattice of six RPCs (Fig S1) [20]. *Drosophila* RPCs (dRPCs) differentiate during the pupal stage and superfluous cells are eliminated by apoptosis [21, 22]. The survival of dRPCs requires cues that are provided by cell-to-cell communication from neighboring cone and primary pigment cells during pupal development [23, 24]. In adult flies, dRPCs have similar functions to mammalian RPCs; that is, they are also juxtaposed to photoreceptor cells and supply metabolites for energy metabolism and other functions [25, 26, 27].

In *Drosophila*, *Fatp* is expressed in both photoreceptors and dRPCs [28], but its physiological role in LD formation in dRPCs and its impact on photoreceptor homeostasis remains to be investigated. Here, we examined the role of FATP and LDs in cellular retinal homeostasis in *Drosophila* and mice. We find that, although the architecture of the mammalian and insect retina is different, RPCs in both organisms play conserved roles in lipid and photoreceptor homeostasis. We show that *Fatp* plays a crucial role in regulating LD production in RPCs and, in turn, RPC-derived LDs have cell non-autonomous effects on the physiology of adjacent photoreceptors.

## Results

### *dFatp* is necessary and sufficient for LD expansion in *Drosophila* retina under physiological and pathological conditions

To investigate the role of *dFatp* in LD formation and expansion, we measured the uptake of BODIPY-C_12_, a fluorescent long-chain FA analog, in the retinas of WT flies (physiological condition) and *Aats-met^FB^* mutants (pathological condition), which accumulate LD due to high mitochondrial ROS levels [9]. BODIPY-C_12_ uptake into LDs (visible as green foci) was increased in the *Aats-met^FB^* mutant retina compared with the WT retina (Fig 1A and 1C). Expression of *dFatp*-specific dsRNA (RNAi) specifically in the RPCs (via the *54C-Gal4* driver) abolished BODIPY-C_12_ uptake in the WT flies and strongly reduced it in the *Aats-met^FB^* mutants (Fig 1A and 1C). We also examined BODIPY-C_12_ uptake in flies with pan-retinal *dFatp* overexpression (using the pan-retinal driver *GMR-Gal4*) and observed a similar accumulation of LDs (Fig 1B and 1C). Importantly, expression of *Brummer*, a gene encoding TAG lipase (referred to here as Bmm-lipase) [29], reduced BODIPY-C_12_ accumulation, confirming that these structures were indeed LDs (Fig 1B and 1C). Collectively, these data demonstrate that *dFatp* is necessary for LD accumulation in the *Drosophila* retina under physiological and pathological conditions.

**Fig 1.**
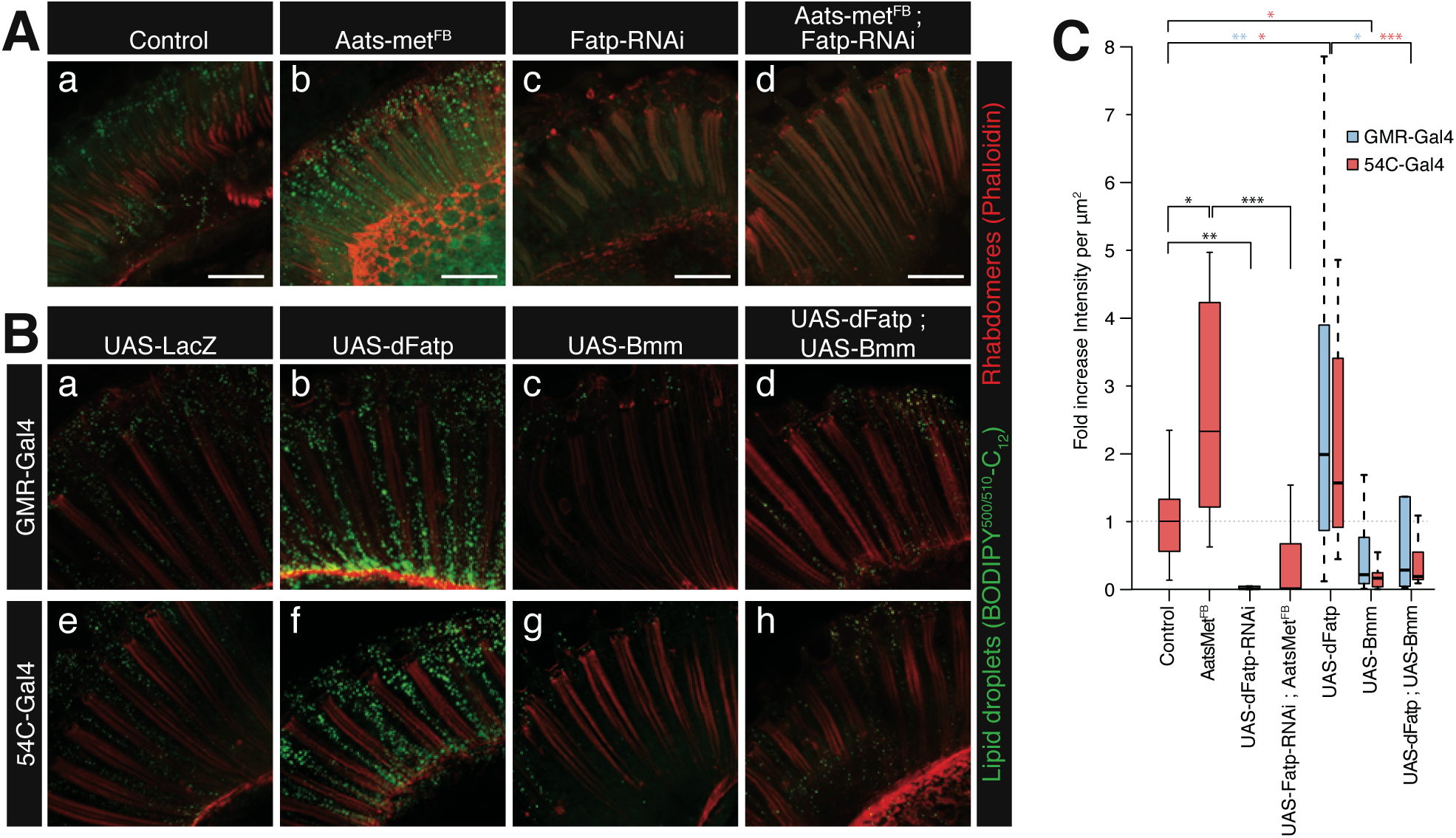
dFatp is required for lipid storage in *Drosophila* retina. (**A**) Confocal fluorescence microscopy of BODIPY-C_12_ (green) uptake in longitudinal sections of whole retinas from one-day-old wild-type (Control) and Aats-met^FB^ mutant flies in the absence (a, b) or presence of dRPC-specific (driven by *54C-Gal4*) dFatp-RNAi^[GD16442]^ (c, d). Photoreceptors were counterstained with phalloidin-rhodamine (red). Scale bar, 25 µm. (**B**) BODIPY-C_12_ (green) uptake in longitudinal sections of retinas from flies expressing *dFatp* and/or *Bmm-lipase* under the control of a pan-retinal (*GMR-Gal4*) (a–d) or a dRPC-specific (*54C-Gal4*) (e–h) driver. Photoreceptors were counterstained with phalloidin-rhodamine (red). Scale bar, 25 µm. (**C**) Quantification of BODIPY-C_12_ uptake into lipid droplets from the images shown in (**A**) and (**B**). Data are presented as the fold change in fluorescence intensity (dots/µm2) compared with the control. The boxes represent the median and lower and upper quartiles, and the whiskers represent the 1.5 interquartile range. N = 6– 41 retinas per condition. Log adjusted values: *p<0.05, **p<0.01, ***p<0.001 by Tukey’s HSD paired sample comparison test.

To confirm these findings and to examine the subcellular localization of LDs in more detail, we performed transmission electron microscopy (TEM) of the retinas of flies with pan-retinal (*GMR-Gal4*) or dRPC-specific (*54C-Gal4*) overexpression of *dFatp* (Fig 2A). LDs were typically visible as homogeneous structures surrounded by a monolayer membrane in dRPCs (Fig 2A and 2B) and, to a much lesser extent, in photoreceptor cells (Fig 2Ac). Interestingly, *dFatp* overexpression significantly increased the size (LD area), but not the number, of LDs in both cell types (Fig 2B and 2C). Finally, dRPC-specific *Bmm-lipase* expression caused a striking reduction in the number of LDs, not only in dRPCs (Fig S2 and 2B) but also in photoreceptors (Fig 2B). This indicates that the monolayer membrane-encapsulated vesicles observed by TEM are indeed LDs, and further suggests that depletion of LDs from dRPCs affects LD homeostasis in neighboring photoreceptor cells.

**Fig 2.**
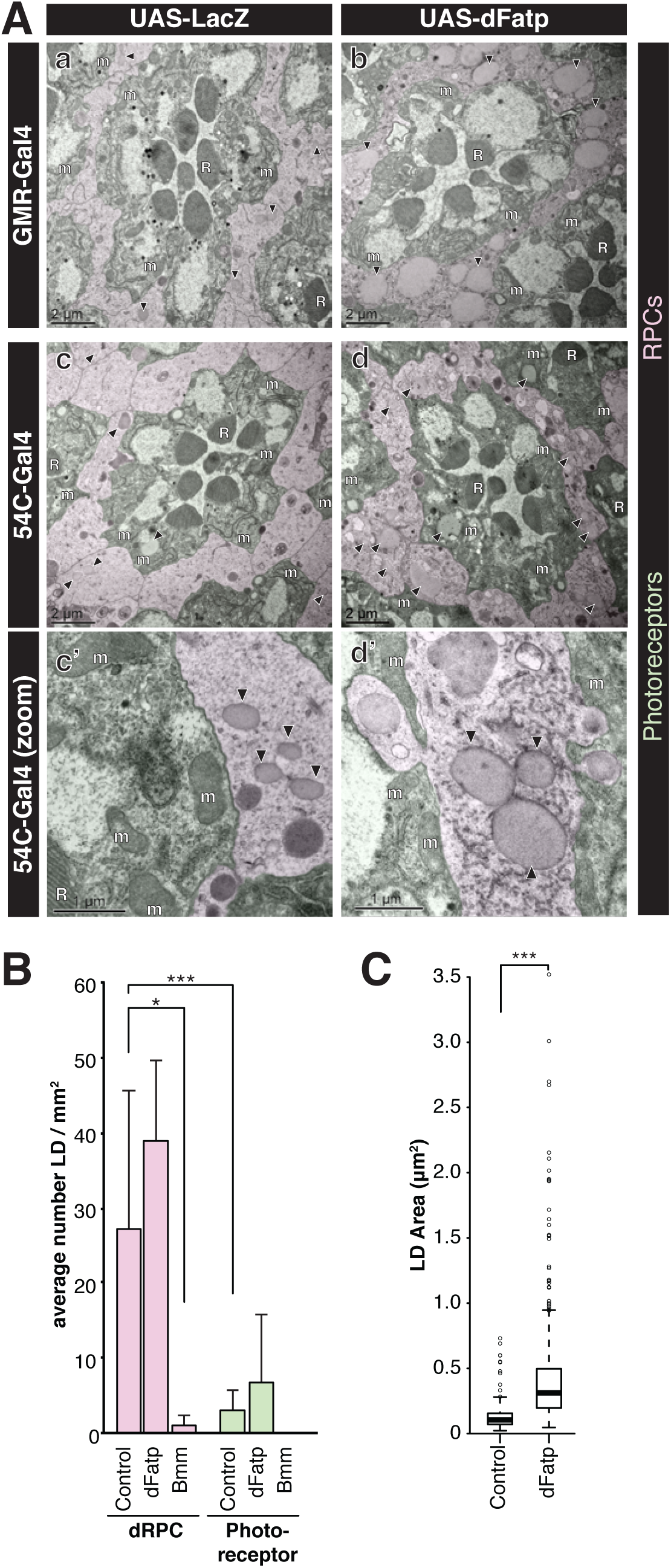
Lipid droplets are mainly localized in the RPCs and are increased by *dFatp* overexpression in *Drosophila* retinas. (**A**) TEM of ommatidia from one-day-old flies expressing *LacZ* (control) or *dFatp* under control of the pan-retinal (*GMR*) or dRPC-specific (*54C*) drivers. One ommatidium in each panel shows seven photoreceptors (false colored green) with central rhabdomeres surrounded by dRPCs (false colored pink). Lipid droplets (black arrowheads) are mainly located in dRPCs. Lipid droplet size was increased by *dFatp* overexpression (b, d, d’). Scale bars, 2 µm (a–d), 1 µm (c’, d’). m, mitochondria; R rhabdomeres; arrowhead, lipid droplet. (**B**) Quantification of lipid droplet density (number per surface area) in dRPCs and photoreceptors in flies with dRPC-specific (*54C*-driven) expression of *LacZ* (control), *dFatp,* or *Bmm-lipase.* Means ± SD of n = 4 eyes. (**C**) Quantification of lipid droplet size in dRPCs of flies with dRPC-specific (*54C*-driven) expression of *LacZ* (control) or *dFatp*. The boxes represent the median and lower and upper quartiles, and the whiskers represent the 1.5 interquartile range. N = 4–5 flies, from which we analyzed >150 fields of view. Log adjusted values: *p<0.05, ***p<0.001 by Tukey’s HSD paired sample comparison tests.

### Cell non-autonomous effects of *dFatp* expressed in dRPCs on photoreceptor LDs and mitochondrial size

The TEM images indicated that dRPC-specific *dFatp* expression increased the size of LDs not only in dRPCs but also in photoreceptors (Fig 2C and Fig 3A) and concomitantly caused a significant non-autonomous decrease in the size of mitochondria in the photoreceptors (Fig 3B). This was confirmed to be a cell non-autonomous effect of dRPCs since co-expression of *dFatp* and *Bmm-lipase* in dRPCs prevented the decrease in photoreceptor mitochondrial size compared with dRPC-specific expression of *dFatp* alone (Fig 3B). Notably, despite the reduction in size, the morphological integrity of mitochondria was preserved, suggesting that their function also remained intact. Nevertheless, the reason for the change in mitochondrial size is unknown.

**Fig 3.**
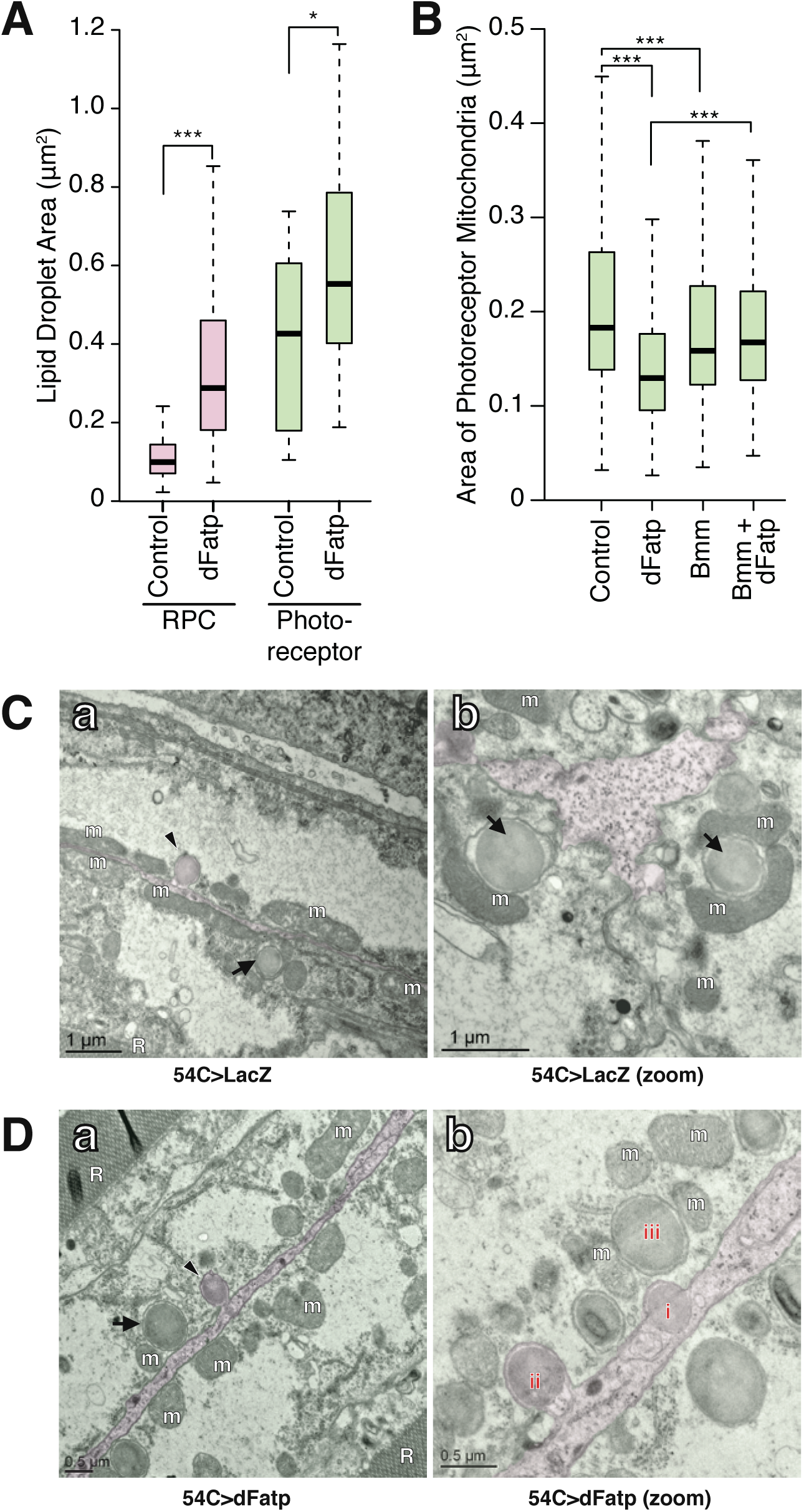
Cell non-autonomous effect of dRPC-specific *dFatp* expression on mitochondrial and lipid droplet size in *Drosophila* photoreceptors. (**A**) Lipid droplet size in dRPCs and photoreceptors of flies with dRPC-specific (*54C*-driven) expression of *LacZ* (control) or *dFatp.* The boxes represent the median and lower and upper quartiles, and the whiskers represent the 1.5 interquartile range. N = 4–5 flies, from which we analyzed >150 fields (calculated from the TEM images in Figure 2). (**B**) Mitochondrial size in photoreceptors of flies with dRPC-specific expression of *LacZ* (control) or *dFatp.* The boxes represent the median and lower and upper quartiles, and the whiskers represent the 1.5 interquartile range. N = 4–5 flies, from which we analyzed >150 fields (calculated from the TEM images in (C) and (D)). *p<0.05, ***p<0.001 by Tukey’s HSD paired sample comparison test. (**C, D**) Low (left) and high (right) magnification horizontal TEM images of retinas of flies with dRPC-specific expression of *LacZ* (**C**) or *dFatp* (**D**). A RPC (false colored pink) is visible sandwiched between two photoreceptors (false colored green). Arrowheads (Ca, Da) indicate lipid droplet-like vesicles entering an endocytotic invagination in the photoreceptor plasma membrane. Arrows indicate granules within photoreceptors surrounded by an additional membrane. (Cb) shows lipid droplets in close association with mitochondria (m) in a photoreceptor. (Db) shows various stages of the vesicle transfer from dRPC to photoreceptor. A vesicle within the dRPC (i) in taken up by the photoreceptor (ii), and surrounded by membrane once internalized within the photoreceptor (iii). Note the position of mitochondria (m) close to the vesicle. Scale bars, 1 µm (C) and 0.5 µm (D).

Interestingly, mitochondria present in photoreceptor cells were preferentially localized close to the membrane adjacent to dRPCs in both WT flies and flies overexpressing dFatp specifically in dRPCs (Fig 3C and 3D). Moreover, the TEM images showed various stages suggestive of the transfer of vesicles from dRPCs to photoreceptors. Thus, in Figure 3Db, vesicles can be seen within the dRPCs (i), in the process of being internalized into the photoreceptor by a process resembling endocytosis (ii), and fully internalized into the photoreceptor, where they appear surrounded by a double membrane (iii). These vesicles were comparable in size to LDs (∼0.25 µm^2^) but appeared slightly more electron dense. Notably, the vesicles were often seen juxtaposed to one or more mitochondria (Fig 3Cb and 3Db). Collectively, these results suggest that *dFatp* expressed in *Drosophila* dRPCs has a non-autonomous effect on adjacent photoreceptors, increasing and decreasing the size of LDs and mitochondria, respectively. While the mechanism by which the information is relayed from dRPCs to photoreceptors is unclear, one possibility is that LDs are physically transferred from dRPCs to photoreceptors, where they become juxtaposed to mitochondria.

### Expression of *hFATP1* in mRPC increases lipid storage and energy metabolism in the mouse retina

To determine whether the function of *dFatp* is conserved, we first asked whether the loss of photoreceptor cells observed in *dFatp^-/-^* mutant retinas [28], could be rescued by the expression of the human homolog *hFATP1* in *Drosophila*. Indeed, photoreceptor-specific expression of *hFATP1* (driven by the rhodopsin 1 [*Rh1*] promoter) strongly reduced the loss of photoreceptors observed in *dFatp^-/-^* mutant retinas (Fig S3). This result supports the conservation of hFATP1 and dFatp function in retinal homeostasis. To investigate the role of hFATP1 in lipid storage in the mammalian retina, we employed our previously described transgenic mice [28], in which hFATP1 is overexpressed using the mammalian mRPC-specific VDM2 promoter (referred to as hFATP1TG mice) [30]. LDs in the retinas of WT (C57BL/6, Fig 4A) and hFATP1TG (Fig 4B) mice were detected by staining of neutral lipids with Nile Red. Notably, transgenic expression of *hFATP1* significantly increased neutral lipid staining, visible as red foci, in the mRPCs of hFATP1TG mice compared with WT mice (Fig 4A–C). Moreover, lipid profiling of dissected mRPCs by gas-chromatography revealed that levels of sterol esters and TAGs, both of which are associated with LDs, were higher in the *hFATP1* mRPCs compared with WT mRPCs (Fig 4D). We also observed that Nile Red staining largely co-localized with perilipin1, a protein associated with LD membranes (Fig 4E–E”), supporting the notion that the Nile Red-stained foci were indeed neutral lipid-containing LDs. Taken together, these data indicate that *hFATP1* expression in mouse RPCs leads to accumulation of LDs, consistent with our findings in *Drosophila* RPCs.

**Fig 4.**
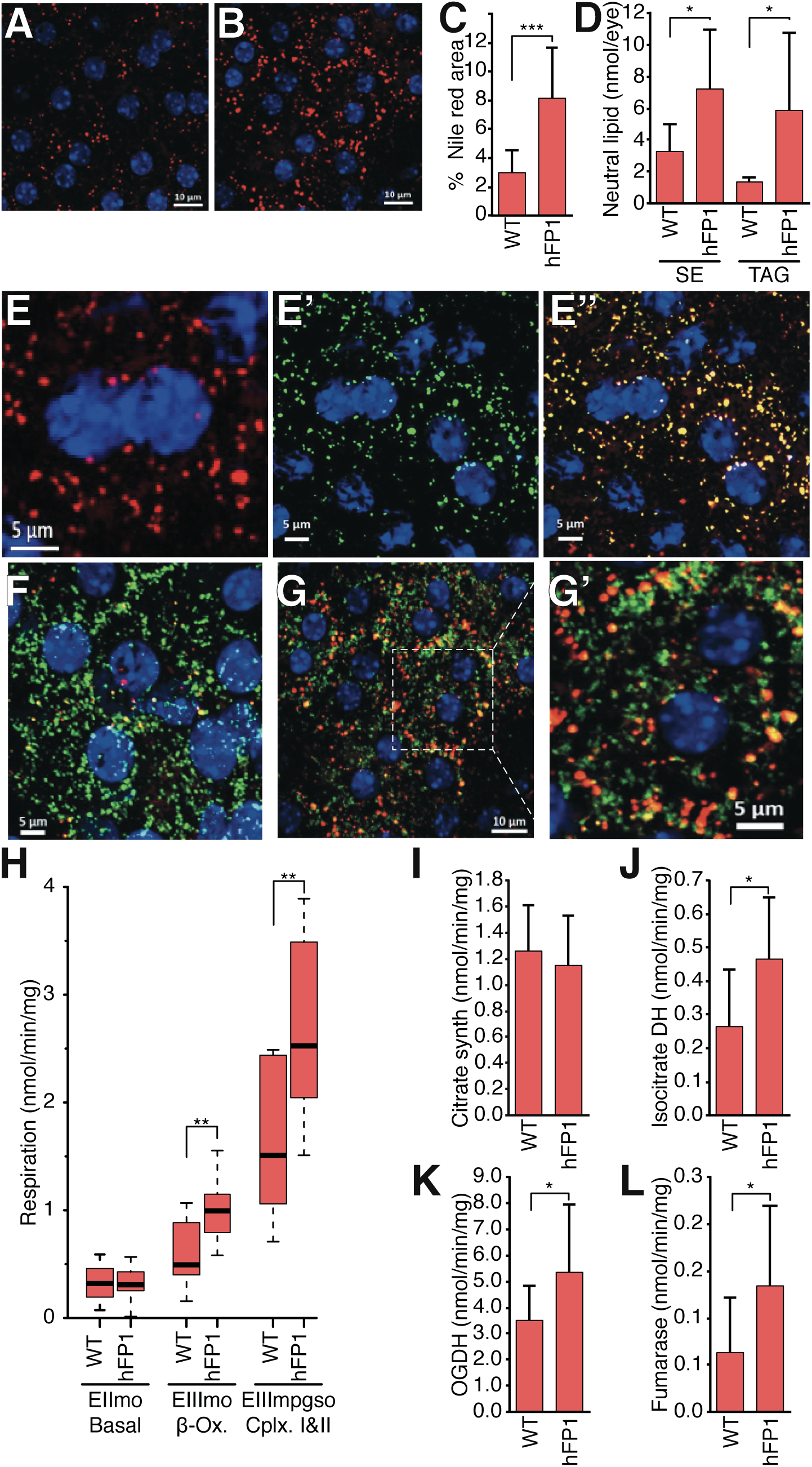
*hFATP1* expression in mouse RPCs increases neutral lipid accumulation and mitochondrial respiration. (**A, B**) Nile Red staining of neutral lipids (red) in mRPCs of a flat-mounted eye cup from wild-type (C57BL/6) mice (**A**) and *hFATP1* transgenic mice (**B**). Nuclei are stained with DAPI (blue). Scale bars, 10 µm. (**C**) Quantification of Nile Red staining in wild-type (WT) and *hFATP1* transgenic (hFP1) mRPCs (as shown in A and B). Mean ± SD of n =19 and 20 animals, respectively. (**D**) Quantification of sterol esters (SE) and triacylglycerides (TAG) in mRPCs from WT and *hFATP1* transgenic mice. Mean ± SD of n = 5 and 7 animals, respectively. (**E–E”**) mRPCs in whole-mount retinas from WT mice double-stained with Nile Red (**E**) and anti-perilipin (green, **E’**). The merged image (**E”**) shows extensive overlap. Scale bars, 5 µm. (**F–G’**) mRPCs in whole-mount retinas of WT (**F**) and *hFATP* transgenic (**G, G’**) mice double-stained with Nile Red and anti-ATP synthase antibody (green, localized to mitochondria). (**G’**) Magnification of the box in (G) shows the juxtaposition of mitochondria and neutral lipid stores. Scale bars, 5 µm (F, G’), 10 µm (G). (**H**) Quantification of mitochondrial respiratory function (O_2_ consumption) in RPCs isolated from WT and *hFATP1* transgenic mice. EIImo, basal respiratory rate; EIIImo, β-oxidation, EIIIMPGSO, global respiratory chain function of mitochondrial complexes I and II. The boxes represent the median and lower and upper quartiles, and the whiskers represent the 1.5 interquartile range. N = 21 and 10 WT and transgenic animals, respectively. (**I–L**) Activities of the TCA cycle enzymes citrate synthase (**I**), isocitrate dehydrogenase (**J**), oxoglutarate dehydrogenase (**K**), and fumarase (**L**) in RPCs isolated from WT and *hFATP1* transgenic mice. Mean ± SD of n > 10 mice for each condition. *p<0.05, **p<0.01, ***p<0.001 for WT vs transgenic groups by two-sample t-test.

Immunofluorescence staining for the mitochondrial marker ATP synthase revealed that LDs and mitochondria were in close proximity in the RPCs of both WT and hFATP1TG mice (Fig 4F–G’), suggesting the possibility that LDs serve as an energy source in mRPCs. To test this hypothesis, we measured the mitochondrial respiratory rate in permeabilized mRPCs by monitoring O_2_ consumption in various mitochondria respiration rates. We found that hFATP1 increased respiration driven by β-oxidation substrates under basal conditions (EIImo basal). mRPCs from hFATP1 transgenic mice also exhibited increased respiration at high steady state rates when respiratory substrates were added (EIIImo β-ox and EIIImpgso CxI+CxII) (Fig 4E). Consistent with this, the activities of the TCA cycle enzymes isocitrate dehydrogenase, 2-oxoglurarate dehydrogenase, and fumarase were all significantly increased in retinal extracts from hFATP1TG mice compared with WT mice (Fig 4J–L). This increased respiration rate in transgenic mRPCs is unlikely to be due to an increase in mitochondrial mass because the activity of citrate synthase was unaffected by hFATP1 overexpression (Fig 4I). Taken together, these data demonstrate that hFATP1 overexpression in mRPCs increases the size of LDs and suggest that they are substrates for energy production in the mitochondria.

Next, we asked whether *hFATP1* expression and enhanced lipid storage in mRPCs could non-autonomously affect the physiology of juxtaposed neural retinal cells, which includes photoreceptor cells, as we had observed in *Drosophila*. mRPCs and the neural retina can be dissociated during dissection, which enables the effects of mRPC-specific *hFATP1* expression on the isolated photoreceptor cells to be examined. We observed an increased number of Oil Red O-stained foci, indicative of neutral lipid-containing droplets, in the neural retina layer of hFATP1TG mice compared with WT mice (Fig 5A–C), suggesting a non-autonomous effect of mRPC-specific *hFATP1* expression similar to the observed phenotype in *Drosophila*. In addition, lipid profiling revealed that sterol ester and TAG levels were higher in the neural retina of transgenic mice compared with WT mice (Fig 5D), and the magnitude of the increase was similar to that seen in isolated mRPCs (compare Fig 5D with Fig 4D). Finally, analyses of the mitochondrial respiratory rate and TCA cycle enzyme activities in neural retinas showed a similar enhancement of O_2_ consumption (Fig 5E) and enzyme activities (Fig 5F–I) in tissue isolated from hFATP1TG mice compared with WT mice.

**Fig 5.**
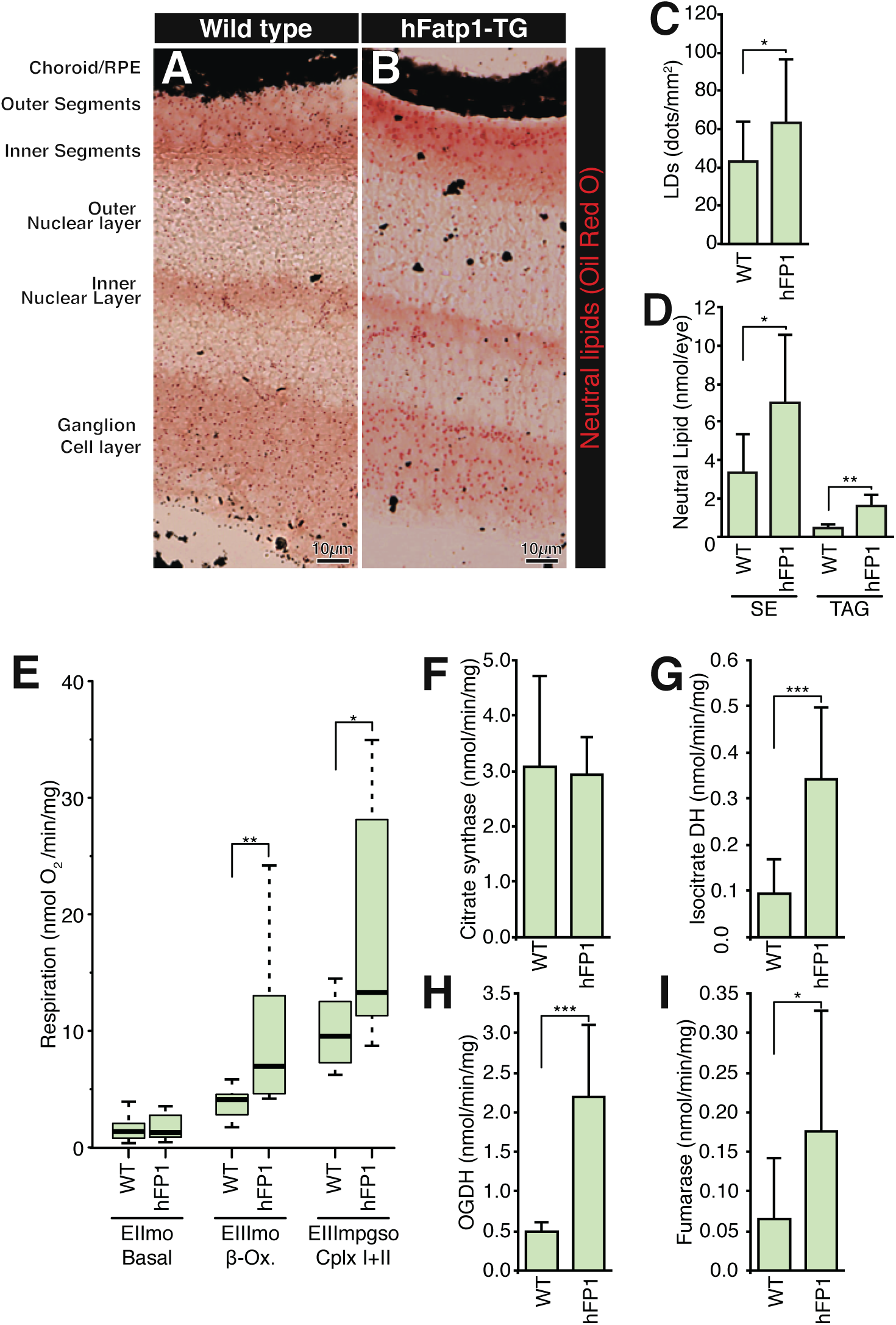
Cell non-autonomous effect of mRPC-specific *hFATP1* overexpression on the lipid content and respiratory rate in the neural retina. (**A, B**) Oil Red O staining of retinal sections of wild-type (WT, **A**) and *hFATP1* transgenic mice (**B**). Scale bars, 10 µm. (C) Quantification of neutral lipid-containing structures seen in (A) and (B). Mean ± SD of n = 10 and 12 WT and transgenic animals, respectively. (**D**) Quantification of sterol esters (SE) and triacylglycerides (TAG) in the neural retina of WT and *hFATP1* transgenic mice. Mean ± SD of n = 5 and 7 retinas, respectively. (**E**) Quantification of mitochondrial respiratory function in the neural retina of WT and *hFATP1* transgenic mice. EIImo, basal respiratory rate; EIIImbgso, β-oxidation; EIIImpgso, global respiratory chain function of complexes I and II. The boxes represent the median and lower and upper quartiles, and the whiskers represent the 1.5 interquartile range. N = 12 and 14 WT and transgenic retinas, respectively. (**F–I**) Activities of citrate synthase (F), isocitrate dehydrogenase (**G**), oxoglutarate dehydrogenase (**H**), and fumarase (**I**) in neural retinal extracts from WT and *hFATP1* transgenic mice. Mean ± SD of n > 9 retinas. *p<0.05, **p<0.01, ***p<0.001 for WT vs transgenic groups by two-sample t-test.

Taken together, these results indicate that *hFATP1* expressed in mRPCs not only increased lipid storage and mitochondrial respiration in the mRPCs themselves, but also in the neural retinal cells. Thus, both the cell autonomous and non-autonomous effects of RPC-expressed FATP1 are conserved in *Drosophila* and mice.

### Dysregulation of LD accumulation perturbs retinal homeostasis

We next examined the consequences of LD accumulation caused by overexpression of *Fatp* for the health of RPCs in *Drosophila* and mice. TEM analyses showed that mRPC-specific expression of *hFATP1* induced vacuolization of the mRPCs and thickening of both mRPC and Bruch’s membranes, which was exacerbated with aging (Fig 6A–D). Similarly, pan-retinal or dRPC-specific expression of *dFatp* caused a loss of dRPCs of about 10%, resulting in perturbation of the organization of ommatidia in *Drosophila* retina (Fig 6E and 6F). In contrast to the dRPCs, photoreceptor cell survival (visualized by expression of Rh1-GFP) was unaffected by *dFatp* overexpression in dRPCs (Fig 6G, left panels, and 6H), which is in agreement with our previous findings in the mouse [30]. Therefore, *Fatp*-mediated LD accumulation in RPCs does not appear to be deleterious to photoreceptor cells under normal physiological conditions. Indeed, dRPC-specific knockdown of *dFatp* resulted in some loss of photoreceptor cells in aged (4-week-old) WT flies (Fig 6G and 6H), suggesting that LDs in RPCs are actually beneficial for photoreceptors during the normal aging process.

**Fig 6.**
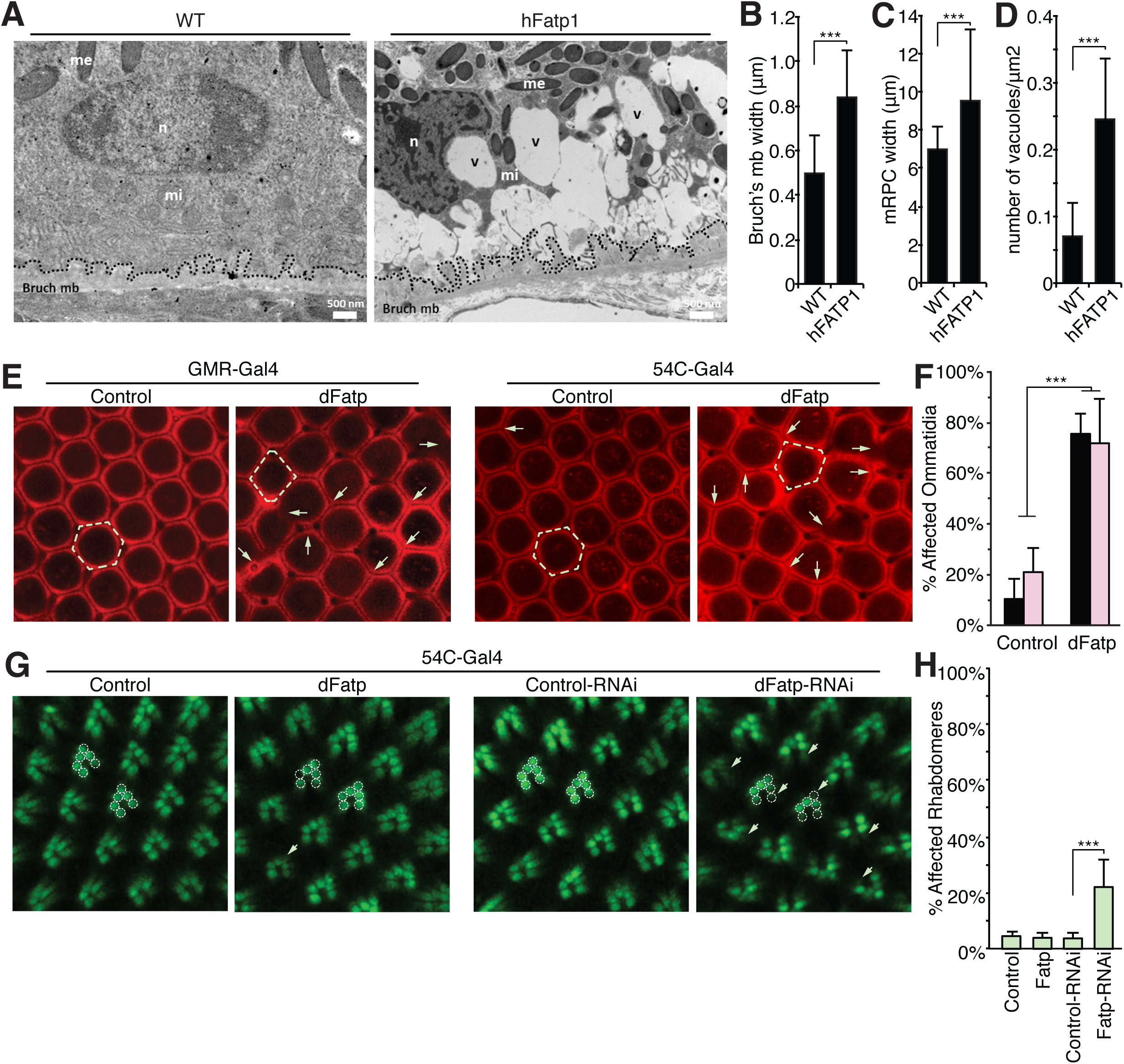
*Fatp* overexpression induces defects in RPCs in both *Drosophila* and mice. (**A**) TEM images of longitudinal sections from 3-month-old wild-type (WT) and *hFATP1* transgenic mice. Abundant vacuoles (v) and thickened Bruch’s membrane (mb, dashed lines) are evident in the aged *hFATP1* mice. me: melanosome, mi: mitochondria, n: nucleus. Scale bars, 500 nm. (**B–D**) Quantification of Bruch’s membrane width (**B**), mRPC width (**C**), and vacuole density in mRPCs (**D**) from the images shown in (A). Mean ± SD of n = 14 and 18 WT and transgenic animals, respectively. (**E**) Confocal fluorescence microscopy (dRPC autofluorescence) of the retinas of 30-day-old *Drosophila* expressing *LacZ* (control) or *dFatp* throughout the retina (*GMR-Gal4*) or in dRPCs alone (*54C-Gal4*). Arrows show loss of dRPCs resulting in disorganization and loss of typical hexagonal ommatidial shape (dashed outline) upon dFatp overexpression. Scale bars, 10 µm. (**F**) Quantification of aberrant ommatidia in retinas shown in (E). Mean ± SD of n = 5 and 8 control and transgenic flies, respectively. Black bars, *GMR-Gal4*; pink bars, *54C-Gal4*. (**G**) Retinas of 40-day-old Rh1-GFP-expressing *Drosophila* with dRPC-specific expression of *LacZ* (control), *dFatp,* control dsRNA (control-RNAi), or *dFatp*-specific dsRNA (*dFatp-* RNAi^[GD9406]^). *dFatp* overexpression in dRPCs does not affect photoreceptor survival, as indicated by intact rhabdomeres (dashed outlines). *dFatp*-RNAi^[GD9406]^ expression induces some loss of rhabdomeres (arrows). Scale bars, 10 µm. (**H**) Quantification of aberrant rhabdomeres, as shown in (G). Mean ± SD of n = 6, 7, and 5 control, *dFatp*-transgenic, and *dFatp*-dsRNA flies, respectively. *p<0.05, **p<0.01, ***p<0.001 by two-sample t-test.

Finally, we asked whether LD accumulation in dRPCs was beneficial or deleterious to photoreceptors under pathological conditions. For this, we examined young (5-day-old) *Aats-met^FB^* mutant flies, in which *dFatp* RNAi prevents LD accumulation at day 1 (Fig 1A and 1C). Importantly, *dFatp* RNAi alone does not induce photoreceptor degeneration in 5-day-old flies [28], but the mutation *Aats-met^FB^* itself causes some degeneration, allowing us to examine the consequence of *dFatp* RNAi in *Aats-met^FB^* mutant on photoreceptor survival. We analyzed the apical/distal position of photoreceptor nuclei, which are normally clustered apically in the WT retina [28] but are mislocalized under conditions associated with photoreceptor death [9]. Monitoring of mislocalized nuclei is a useful readout of cell death in situations where photoreceptor rhabdomeres are difficult to count due to their rapid degradation, as is the case for *Aats-met^FB^* mutants. We observed an increase in the number of nuclei mislocalized between the proximal and distal part of the retina in *Aats-^B^* mutants compared with WT flies, consistent with previous reports for this mutant [9]. However, the mislocalization was completely reversed by dRPC-specific *dFatp* RNAi (Fig 7).

**Fig 7.**
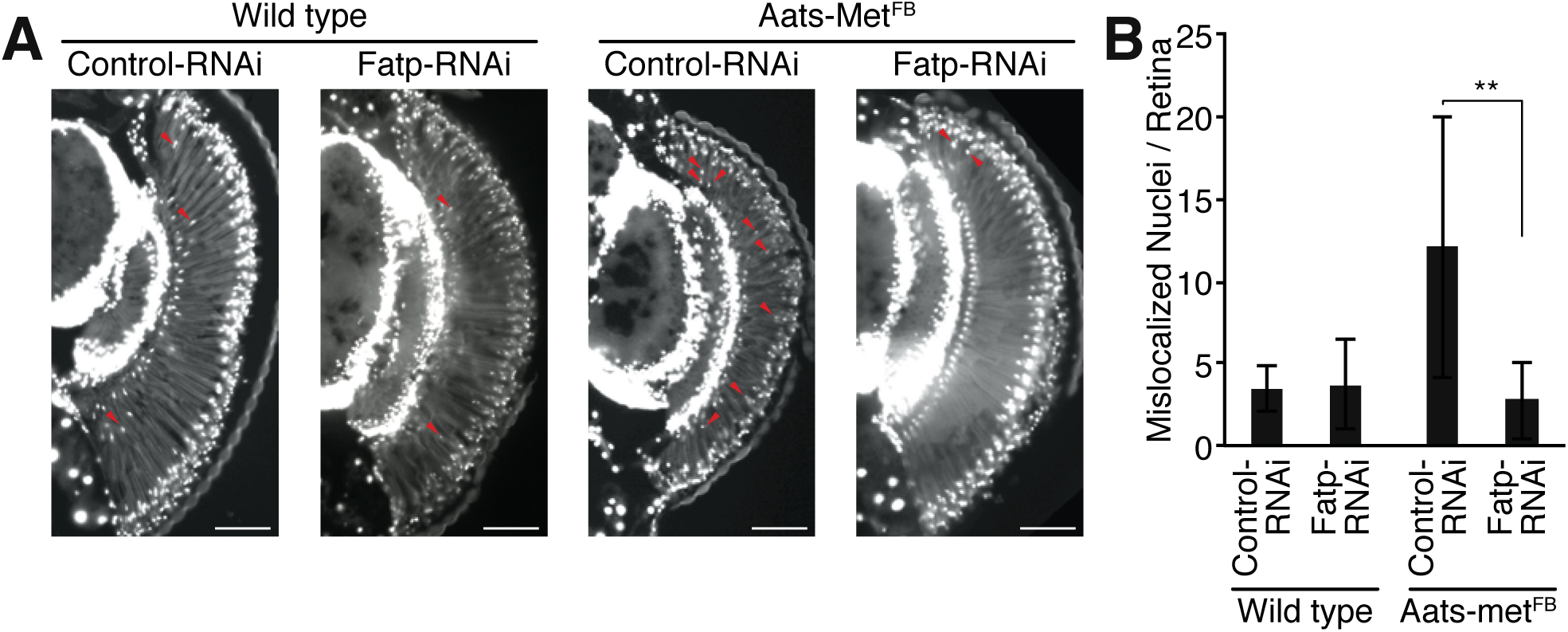
dRPC-specific RNAi of *Fatp* suppresses photoreceptor degeneration in *Drosophila* retina. (**A**) Epifluorescence microscopy of retinal sections from 5-day-old wild-type or *Aats-met^FB^* mutant flies with or without Fatp-RNAi^[GD16442]^. In the mutant, nuclei (stained with DAPI, white) are mislocalized between the distal and proximal part of the retina (red arrows), indicative of dying photoreceptors. dRPC-specific expression of Fatp-RNAi^[GD16442]^ (*54C-Gal4* drive) markedly reduces the number of dying photoreceptors. Scale bars, 100 µm. (**B**) Quantification of mislocalized nuclei, as shown in A. Mean ± SD of n > 6 retinas.**p<0.01 by two-sample t-test.

Altogether, these results indicate that Fatp-mediated LD accumulation in dRPCs is harmful to photoreceptors under conditions of pathological oxidative stress (*Aats-met^FB^* mutants), in stark contrast to the apparent beneficial effect under physiological conditions.

## Discussion

Here, we studied the consequences of dysregulated lipid storage in RPCs on photoreceptor structure and function in both *Drosophila* and mouse retinas. Our results show that the mechanisms of lipid storage and communication between RPC and photoreceptor layers are conserved in flies and mice. Thus, we propose the *Drosophila* RPC–photoreceptor interaction as a novel paradigm for modeling the relationship between mammalian RPCs and photoreceptors under physiological and pathological conditions.

We made several novel observations in this study. *FATP* expression is required for LD formation in RPCs. FATP has acyl-CoA synthetase activity that is thought to facilitate cellular uptake of FAs (reviewed in [16]). However, FATP also interacts with DGAT-2 on the endoplasmic reticulum, where it has been show to function in LD biogenesis in *C. elegans* [13]. Interestingly, LD accumulation induced by *FATP* overexpression resembles one of the key hallmarks of AMD pathology; that is, the accumulation of lipids in RPCs and of drusen in Bruch’s membrane [31]. We also found that *FATP* expression in RPCs induced cell non-autonomous accumulation of LDs in the adjacent photoreceptor layer. In mice, mRPC-specific *hFATP1* expression also increased β-oxidation, TCA enzyme activity, and mitochondrial respiration in photoreceptors. We did not evaluate the mitochondrial respiration rate in the *Drosophila* model due to the technical difficulty of dissociating RPCs from photoreceptor cells. However, the mitochondrial architecture in photoreceptor cells was normal by TEM. LDs were frequently observed juxtaposed to mitochondria in both flies and mice, suggesting that lipase-mediated release of TAGs from LDs provides an energy source for the mitochondria. This is supported by studies showing that physical proximity between LDs and mitochondria is important for β-oxidation of FAs [32]. Thus, both the function of FATP in RPCs and its effect on photoreceptor cells are conserved from flies to mammals. Despite the difference in mouse and fly retinal architecture, RPCs appear to be functionally homologous in both species with respect to providing metabolic support for adjacent photoreceptors. *Drosophila* RPCs form a tight barrier between the hemolymph and photoreceptors, similar to the mammalian outer retina [33], implying that dRPCs, like mammalian RPCs, may play an active role in nutrient transport [31]. We suggest that *Drosophila* may be a useful model to investigate mammalian RPC disorders, such as AMD, particularly given its genetic tractability.

In our previous work, we showed that *dFatp* is expressed at higher levels in dRPCs than in photoreceptors, and that loss of dFatp in both cell types led to photoreceptor degeneration [28]. We had proposed that degeneration of photoreceptors in the dFatp mutant is due to a failure to degrade Rh1, which accumulates to toxic levels and triggers apoptosis. We now show that dFatp expressed in dRPCs also contributes to photoreceptor viability via its role in LD biosynthesis. Therefore, dFatp appears to have two distinct functions in photoreceptors and dRPCs. In photoreceptors, dFatp is required for optimal Rh1 metabolism, presumably due to its role in phosphatidic acid synthesis and Rh1 trafficking [16, 34, 35]. In RPCs, dFatp is required for expansion of LDs, which can then be metabolized for energy production. The loss of photoreceptors following RPC-specific depletion of *dFatp* may therefore be due to the dwindling supply of energy, especially since photoreceptors have high energy demands [5]. This hypothesis is in agreement with recent reports demonstrating the importance of *de novo* LD biogenesis in protecting photoreceptors against lipotoxicity under conditions of low nutrient supply or high energy demand [36, 37]. Finally, hFatp1 has been shown to directly interact with visual cycle enzymes [38.] Expression of human *Fatp1* in mouse RPCs leads to accumulation of retinoids, which are nontoxic to photoreceptors except under conditions of excessive light exposure [30]. Whether or not loss or overexpression of *dFatp* leads to dysregulation of the visual cycle in flies remains to be established.

Our data establish that LDs are not toxic to retinal cells under physiological conditions, but that they can contribute to neurodegeneration under some pathological conditions—as shown here with the *Aats-met^FB^* mutants (model in Fig 8). In these mutants, suppression of LD accumulation in dRPCs is neuroprotective, suggesting that the LDs non-autonomously promote photoreceptor cell death under conditions of oxidative stress. This is consistent with studies of flies with mitochondrial defects that cause high ROS levels in neurons, which demonstrated that removal of LDs by *dFatp* knockdown rescued retinal degeneration in *sicily* (*Drosophila* homolog of the human nuclear encoded mitochondrial gene NDUFAF6) and *marf* (*Drosophila* homolog of the mitochondrial fusion GTPases, Mitofusin 1 and 2) mutants and that ectopic expression of *Bmm-lipase* rescued photoreceptors in *Aats-met* mutants [9, 27]. One possible explanation for the deleterious effects of LDs in these mutants is that LD-mediated enhancement of mitochondrial respiration may have resulted in toxic levels of ROS and/or lipid peroxidation, leading to cell death.

**Fig 8.**
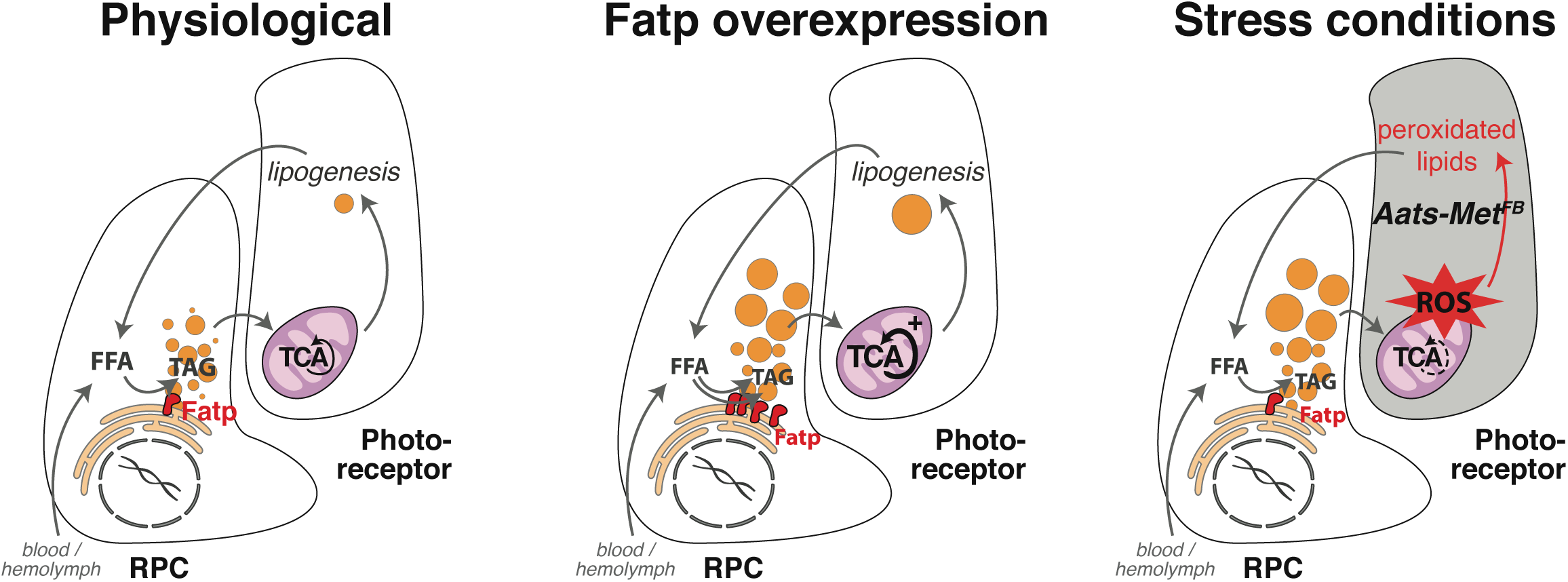
Schematic of the role of FATP in the metabolism of lipid droplets and communication between RPCs and photoreceptors under physiological and pathological conditions. Under physiological conditions, FATP is involved in the maintenance of lipid droplets in RPCs. Overexpression of FATP in RPCs expands the cellular lipid droplet content, which has a cell non-autonomous stimulatory effect on neighboring photoreceptors (possibly by direct intercellular transfer of lipid vesicles) that increases mitochondrial oxidation. This is observed in *Drosophila* and mice. Under conditions of oxidative stress (e.g., *Aats-met^FB^* mutant flies, in which ROS levels are elevated in photoreceptors), the dRPC-signaled boost in mitochondrial oxidation is deleterious to photoreceptors, leading to dysfunction and death.

We found that LD accumulation in RPCs induced by ectopic expression of *FATP* is not toxic to photoreceptors in either *Drosophila* (this study) or mice [30], suggesting that LDs are not deleterious under normal physiological conditions. In addition to acting as an energy source, LDs may also reduce the cellular sensitivity to ROS. The latter possibility is supported by the finding that light exposure, which induces the accumulation of photodamaged proteins and lipids [39], enhances photoreceptor degeneration to a greater extent in *dFatp* mutant flies than in WT flies [28]. Similarly, LD accumulation in glia protects against ROS-induced damage of the developing nervous system of *Drosophila* upon exposure to hypoxia [8]. Moreover, LDs are not uniform structures but exist in various forms with differing potential functions, such as storage of phospholipids, vitamin E, and cellular toxins [10]. Taken together, our findings and those of others suggest that LDs may play a protective or beneficial role under physiological or moderately stressful conditions.

Although our data demonstrate that FATP-mediated LD accumulation in RPCs provides a metabolic signal to photoreceptors, it is not clear whether the signal itself is a lipid or another signaling molecule. Recent work proposed that in mitochondrial respiratory chain-deficient mutants, the accumulation of LDs in RPCs induces transfer of lactate from RPCs to photoreceptors [27]. The increased lactate levels in photoreceptors enhanced the activity of the TCA cycle and synthesis of FAs, which then underwent apolipoprotein-mediated transfer back to RPCs and induced LD formation [27]. A similar mechanism was described in mouse neuronal/glial co-cultures, raising the possibility that it may also be operating in the retina of hFATP1TG mice [27]. Our ultrastructural analysis supports the possibility of an exchange of electron dense material between RPCs and photoreceptor cells in *Drosophila*. Indeed, electron dense vesicles—similar in appearance and size to LDs—were observed in contact with dRPCs, as invaginations in the photoreceptor membrane, and finally as double-membraned structures in close proximity to the mitochondria in photoreceptors. Associations between LDs and mitochondria have also been observed in skeletal and heart muscle, where it was proposed to facilitate β-oxidation [40, 41, 32]. Although further studies will be required, our ultrastructural findings provide a hint that vesicle transfer could be a route of communication between RPCs and photoreceptors. Collectively, our results demonstrate that crosstalk between RPCs and photoreceptors is crucial for normal photoreceptor homeostasis and that its breakdown may contribute to retinal pathologies.

## Material and Methods

### Drosophila genetics

The fly stocks used in this study were: *54C-Gal4* (BS#27328) [24], *GMR-Gal4*, *Rh1-Gal4* [42], *UAS-dFatp, FRT40A Fatp^k10307^* [28], *UAS-bmm* [29], *UAS-dFatp-RNAi* (VDRC stocks GD16442, GD9604 and KK100124), *FRT82^B^ Aats-met^FB^* (BS#39747), and *Rh1-GFP* [43]. Mitotic whole-eye clones for *Fatp^k10307^* and *Aats-met^FB^* were generated by crossing a mutant line carrying the FRT-containing chromosome with a line containing *ey-flp* and *FRT40A GMR-hid* (for Fatp^k10307^) or *FRT82B GMR-hid* (for *Aats-met^FB^*) while mosaic eye clones were made using the line *Rh1-Gal4,ey-flp;FRT40A Rh1-TdTomato^NinaC^;UAS-GFP* crossed to *FRT40A Fatp^k10307^;UAS-hFatp1* [44, 45]. Human Fatp1 cDNA, a gift from Celine Haby (IGBMC Strasbourg), was cloned via BamHI and EcoRI into a pUAST-w+-attB transgenic vector. Transgenic lines were generated by Best Gene (Chino Hills, CA, USA) using PhiC31 integrase-mediated transgenesis [46] at the same site used for dFatp (65B2). Flies were maintained on standard corn medium at 25°C.

### Lipid droplet staining

The retinas of 1-day-old flies were dissected in ice-cold HL3 medium [47] according to [28]. Tissues were warmed to 25°C and incubated for 30 min with 1.5 µg/mL C_1_-Bodipy-500/510-C_12_ (BODIPY-C_12_; D3823, Life Technologies). Samples were washed three times in HL3 medium, fixed in 4% paraformaldehyde (PFA, Electron Microscopy Sciences) for 15 min at room temperature (RT), washed three times in wash buffer (PBS containing 0.1% Triton-X100), and incubated for 16 h at 4°C in wash buffer supplemented with 25 ng/mL phalloidin-TRITC (Sigma). The samples were washed three times in wash buffer, mounted on a bridge slide in Vectashield (Vector Laboratories), and stored at −20°C until analysis. 16-bit image stacks were acquired on a Zeiss LSM800 microscope and processed for quantification using ImageJ2 software [48]. Images were filtered for noise using Gaussian Blur 3D (σ = 1), after which a Z-projection was made. LDs were identified by thresholding the images, and the integrated density of the signal and total retinal area were measured. The integrated density of BODIPY staining divided by the total area of the retina was normalized to the control for each experiment.

### Statistical analysis

Statistical analyses were performed using R. Group differences were analyzed by t-test or Tukey’s HSD paired sample comparison test, as specified in the figure legends.

### Imaging of photoreceptors and RPCs in living flies

Living flies, maintained for 30 days at 25°C on a 12h light/dark cycle, were anesthetized using CO_2_, embedded in 1% agarose covered with cold water [45], and imaged using a Leica SP5 upright confocal microscope. RPCs were visualized by pigment autofluorescence (excitation 514 nm, detection 530–630 nm) and photoreceptors were identified by Rh1-GFP expression (excitation 488 nm, detection 500–570 nm). RPCs and photoreceptor rhabdomeres were quantified using Fiji cell counter tool.

### Transmission electron microscopy

Dissected *Drosophila* eyes were fixed in 0.1 M cacodylate buffer, 2.5% glutaraldehyde, and 2 mM CaCl_2_ for 16 h at 4°C. After rinsing with 0.1 M cacodylate buffer at RT, the eyes were incubated with 1% OsO_4_ in 0.1 M cacodylate buffer for 2 h at RT. Tissues were then progressively dehydrated in acetone at RT and mounted in 100% epoxy resin (Epon 812) in silicone-embedding molds. After resin polymerization for 48 h at 60°C, samples were sliced into 60 nm sections, which were stained with lead citrate and examined with a Philips CM120 transmission electron microscope (TEM) operating at 80 kV.

### Mice

Transgenic mice overexpressing human *Fatp1* (hFATP1TG) specifically in the RPCs (driven by the RPC-specific VDM2 promoter) were generated on a C57BL/6J genetic background as previously described [30]. The hFATP1TG mice were maintained on a standard 12 h light (90 lux)/12 h dark cycle at ∼22°C and were fed *ad libitum* with a standard rodent diet. Mice were housed in facilities accredited by the French Ministry of Agriculture and Forestry (B-34 172 36—March 11, 2010). Experiments were carried out in accordance with the European Communities Council Directive of 24 November 1986 (86/609/EEC) and the French Ethical Committee (CEEA-LR-12141) guidelines for the care and use of animals for experimental procedures. Mice were euthanized by cervical dislocation and the eyes were enucleated and dissected.

### Co-staining of lipids, perilipin, and ATP synthase

Nile Red staining of lipids [49] was used alone or in conjunction with fluorescent protein immunostaining. The eyes of 3-month-old mice were enucleated and the retina was separated from the RPC/choroid layer to obtain an empty eyeball [30]. The eyes were fixed in 4% PFA for 1 h at RT, washed with PBS, and permeabilized with 0.1% sodium dodecyl sulfate. They were then incubated for 20 min at RT in blocking buffer (10% fetal calf serum in PBS), and incubated overnight at 4°C with a mouse anti-mouse ATP synthase (Millipore MAB3494, 1:500) or rabbit anti-mouse perilipin (D1D8, Cell Signaling Technology # 9349, 1: 200) primary antibody. The tissues were then incubated for 4 h at RT with Alexa Fluor 488-conjugated anti-rabbit or anti-mouse secondary antibodies diluted in blocking buffer. The immunostained eyeballs were gently rinsed in PBS, incubated with Nile Red solution (10 µg/ml) for 30 min at RT in the dark, washed twice in PBS for 5 min at RT, and incubated with 4′,6-diamidino-2-phenylindole (DAPI, 1:1000) for 5 min at RT. The eyeballs were finally rinsed 5 times in PBS for 5 min at RT and mounted in Dako mounting medium. Confocal imaging was performed with a Zeiss LSM 5 LIVE DUO Highspeed/Spectral Confocal system. Images were acquired with Zeiss Zen software, and LDs were counted with ImageJ software.

### Oil Red O staining

Neutral lipids in frozen sections of the retina were stained with Oil Red O. Mouse eyes were rapidly enucleated and fixed in 4% PFA for 4 h at 4°C. Whole eyeballs were embedded in OCT compound (Tissue Tek #4583) and cut into 10 µm sagittal sections. The sections were thawed at RT for 10 min, incubated in PBS for 20 min at RT, and then incubated in a freshly prepared Oil Red O (Sigma-Aldrich # O0625) working solution (3 ml of 5 mg/mL Oil Red O in isopropanol mixed with 2 ml of H_2_O and filtered through 0.4 µm filter) for 30 min in the dark. The sections were washed 5 times with water, mounted in Mountex medium, and visualized with a NanoZoomer 2.0–HT digital scanner (Hamamatsu).

### Quantification of neutral lipids in mouse retina

Mice were euthanized by vertebral dislocation, the eyes were enucleated, and the neural retina was removed from the eye cup. Neural retinas were directly frozen in liquid nitrogen. The RPC/choroid was scraped and collected in PBS, and the samples were centrifuged to remove the PBS. Tissues were stored dry at −80°C.

Lipids were extracted and analyzed as previously described [50]. Total lipids were extracted twice from tissues with ethanol/chloroform (1:2, v/v). Before extraction, internal standards were added. The organic phases were dried under nitrogen and lipid classes were separated by thin-layer chromatography on silica gel G using a mixture of hexane-ethyl ether-acetic acid (80:20:1 v/v/v). Lipids were transmethylated and the fatty acid methylesters were analyzed by gas chromatography. Briefly, samples were treated with toluene-methanol (1:1, v/v) and boron trifluoride in methanol. Transmethylation was carried out at 100°C for 90 min in screw-capped tubes. After addition of 1.5 mL, 10% K_2_CO_3_ in water, the resulting fatty acid methyl esters were extracted with 2 mL of isooctane and analyzed by gas chromatography, using an HP6890 instrument equipped with a fused silica capillary BPX70 SGE column (60 × 0.25 mm). The vector gas was hydrogen. The temperatures of the Ross injector and the flame ionization detector were set at 230°C and 250°C, respectively

### Oxygen consumption

Respiration was measured on RPC/choroid and neural retinas permeabilized by incubation for 2 min with 15 µg digitonin per mg and resuspended in a respiratory buffer (pH 7.4, 10 mM KH_2_PO_4_, 300 mM mannitol, 10 mM KCl and 5 mM MgCl_2_). The respiratory rates were recorded at 37°C in 2 ml glass chambers using a high-resolution Oxygraph respirometer (Oroboros, Innsbruck, Austria). Assays were initiated in the presence of 5 mM malate/0.2 mM octanoyl carnitine to measure state 2, basal respiration (EIImo basal). Complex I-coupled state 3 respiration was measured by adding 0.5 mM NAD^+^/1.5 mM ADP (EIIImo β-ox). Then, 5 mM pyruvate and 10 mM succinate were added to reach maximal coupled respiration (EIIImpgso CxI+CxII), and 10 µM rotenone was injected to obtain the CII-coupled state 3 respiration. Oligomycin (8 µg/mL) was added to determine the uncoupled state 4 respiration rate. Finally, carbonyl cyanide-4-(trifluoromethoxy) phenylhydrazone (1 µM) was added to control the permeabilization of the tissues.

### Mitochondrial enzymatic activities

The activities of the mitochondrial citrate synthase, oxoglutarate dehydrogenase, isocitrate dehydrogenase and fumarase were measured in RPC/choroid and neural retina homogenates at 37°C using a Beckman DU-640B spectrophotometer (Beckman Coulter) or a CLARIOstar (BMG LabTech) [51]. Briefly, citrate synthase activity was measured using 0.15 mM DTNB reagent (SIGMA Aldrich) which interacts with CoA-SH to produce TNB. The formation of TNB was followed for 1.5 minutes at wavelength of 412 nm. IDH and OGDH activities were determined in the presence of respective substrates isocitrate and α-ketoglutarate, by monitoring for 3 minutes the change in NAD^+^ to NADH which absorbs light at 340 nm. Fumarase activity was determined by measuring the conversion of L-malate to fumarate, monitoring the increase in absorbance at 250 nm. The optical density variation per minute is calculated from the curve and the enzymatic activity expressed as nmol of product formed / minute / mg protein.

### Electron microscopy of mouse RPCs

The eyes of 3-6 month old hFATP1TG mice were rapidly enucleated, the corneas were removed, and the eyeballs were fixed by immersion in 2.5% glutaraldehyde in Sorensen’s phosphate buffer (0.1 M, pH 7.4) overnight at 4°C. The tissues were then rinsed in Sorensen’s buffer and post-fixed in 1% OsO_4_ for 2 h in the dark at RT. The tissues were rinsed twice, dehydrated in a graded series of ethanol solutions (30–100%), and embedded in EmBed 812 using a Leica EM AMW (Automated Microwave Tissue Processor for Electronic Microscopy). Sections (60 nm thick) were cut near the optic nerve (Leica-Reichert Ultracut E), counterstained with uranyl acetate, and examined using a Hitachi 7100 TEM (Centre de Ressources en Imagerie Cellulaire de Montpellier, France). The thickness of Bruch’s membrane was determined by measuring both the thickest and the thinnest parts of 5 fields throughout the retinal section per mouse. Data are presented as the median value per eye. We also enumerated the vacuoles in 5 fields of retinal sections per mouse at 10,000× magnification.

## Acknowledgements

We are grateful to the ARTHRO-TOOLS and the PLATIM microscopy platform of SFR Biosciences (UMS3444/CNRS, US8/INSERM, ENS de Lyon, UCBL) and the Centre Technologique des Microstructures CTµ at Lyon1 for assistance with electron microscopy. We are grateful to the assistance of Victor Girard in preparing the schematic for Fig 8. Stocks were obtained from the Bloomington Drosophila Stock Center (NIH P40OD018537). This work was supported by the French National Research Agency award ANR-12-BSV1-0019-02 to BM, PB, DVDB, and AC and by a postdoctoral fellowship to FN from the Association Française contre les Myopathies, a European Union’s Seventh Framework Programme/AIRC (Associazione Italiana per la Ricerca sul cancro) Reintegration Grant.

