## Supplementary Materials for "Expression of fatty acid transport protein in retinal pigment cells promotes lipid droplet expansion and photoreceptor homeostasis"

**Supplemental Figure legends.**

**S1 Fig. Schematic of the mouse and *Drosophila* retina.**

(**A**) Mouse retinal pigment epithelial cells (mRPCs, pink) support photoreceptor rods and cones (green) by providing them with nutrients, transported across Bruch's membrane from the underlying vasculature. (**B**) Horizontal and tangential sections through a *Drosophila* ommatidium (of ~800 in total) showing dRPC organization around the photoreceptors.

**S2 Fig. dRPC-specific *Bmm-lipase* expression reduces lipid droplet content.**

Transmission electron microscopy of a single ommatidium showing photoreceptors (false colored green) and dRPC (pink). (**A**) Lipid droplets (e.g., black arrowhead) are reduced in number and appear as empty vesicular structures in (**A, A'**, open arrowheads). Scale bars, 2 µm (A), 1 µm (A'). R, rhabdomeres; m, mitochondria.

**S3 Fig. dFatp is a functional ortholog of human FATP1.**

(**A, B**) Confocal fluorescence microscopy of retinas from *dFatp*$^{-/-}$ (*dFatp*$^{k10307}$) mutant flies without (A) or with (B) photoreceptor-specific expression of human *FATP1* (*dFatp*$^{-/-}$ *;Rh1>hFATP1*). *hFATP1* rescues the loss of photoreceptors in *dFatp*$^{-/-}$ mutant clones. Retinas were analyzed using the Tomato/GFP-FLP/FRT technique, in which all photoreceptors are marked by GFP, and homozygous mutant mosaic retina is marked by the absence of TdTomato. *dFatp*$^{k10307}$-mutant tissue shows loss of photoreceptors at 15 days of age (**A**). Scale bars, 10 µm. (**C**) Quantification of photoreceptor loss, as shown in (A) and (B). Mean ± SD of n = 12 retinas. ***p<0.001 by two-sample t-test.

### Figure S1

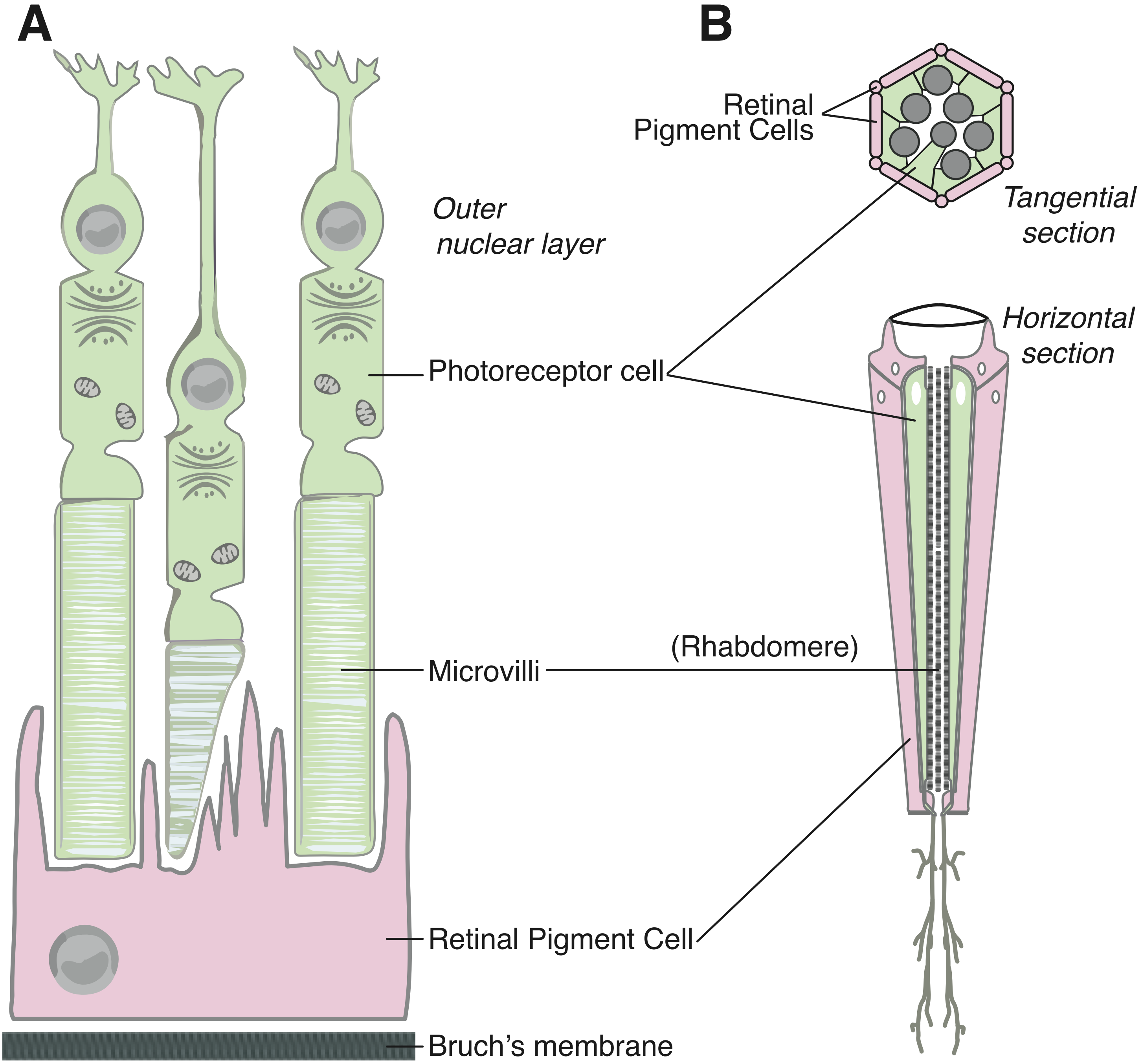
A
B
Outer
nuclear layer
Photoreceptor cell
Microvilli
Retinal Pigment Cell
Bruch's membrane
Retinal
Pigment Cells
Tangential
section
Horizontal
section
(Rhabdomere)

### Figure S2

A 54C>Bmm A’ 54C>Bmm

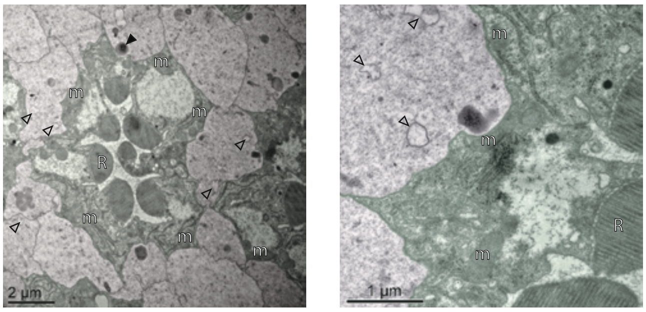

### Figure S3

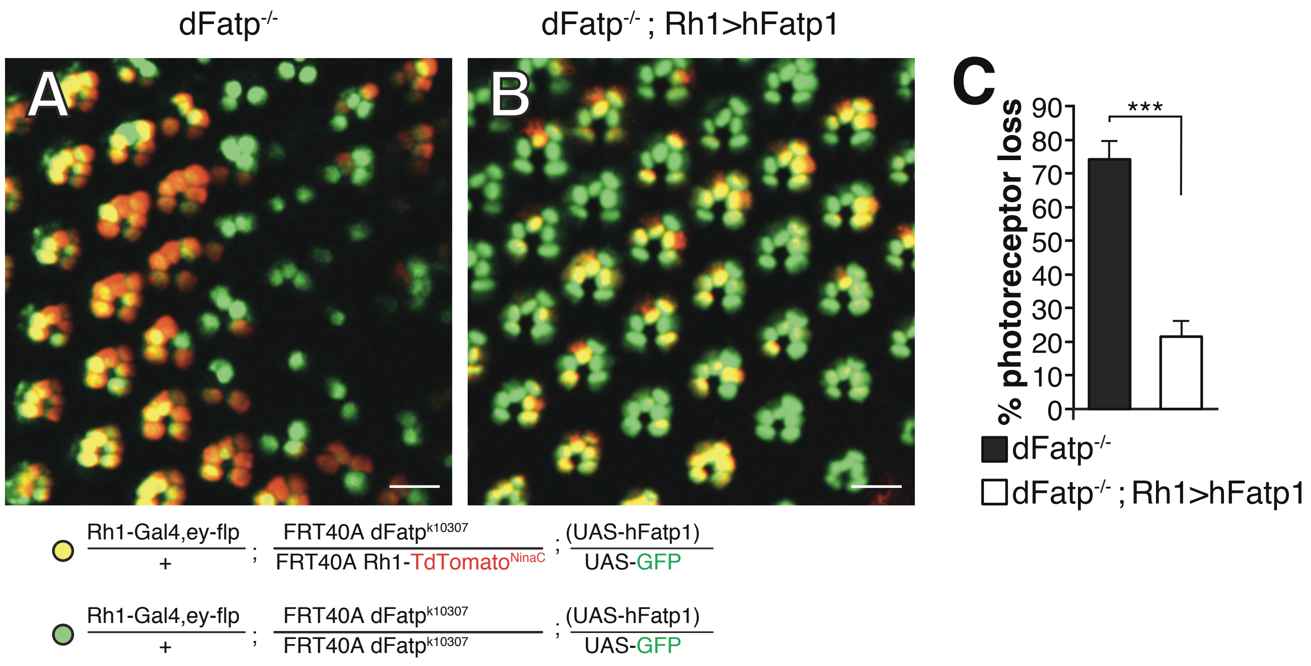
dFatp-/-
dFatp-/- ; Rh1>hFatp1
A
B
C
% photoreceptor loss
0
10
20
30
40
50
60
70
80
90
***
dFatp-/-
dFatp-/- ; Rh1>hFatp1
Rh1-Gal4,ey-flp / + ; FRT40A dFatpk10307 / FRT40A Rh1-TdTomatoNinaC ; (UAS-hFatp1) / UAS-GFP
Rh1-Gal4,ey-flp / + ; FRT40A dFatpk10307 / FRT40A dFatpk10307 ; (UAS-hFatp1) / UAS-GFP
